# But will they even *wear* it? Exploring the tolerability of social communication coaching smartglasses in children and adults with autism

**DOI:** 10.1101/164376

**Authors:** Neha U. Keshav, Joseph P. Salisbury, Arshya Vahabzadeh, Ned T. Sahin

**Affiliations:** Brain Power, LLC, Cambridge, MA, United States; Department of Psychiatry, Massachusetts General Hospital, Boston, MA, United States; Department of Psychology, Harvard University, Cambridge, MA, United States

## Abstract

**Introduction:** Augmented reality smartglasses are an emerging technology that are under investigation as a social communication aid for children and adults with autism spectrum disorder (ASD), and as a research tool to aid with digital phenotyping. Tolerability of this wearable technology in people with ASD is an important area for research, especially as these individuals may experience sensory, cognitive, and attentional challenges.

**Aims:** The aim of this study was to assess the tolerability and usability of a novel smartglasses system that has been designed as a social communication aid for children and adults with autism (the *Brain Power Autism System;* BPAS). BPAS runs on Google Glass Explorer Edition and other smartglasses, utilizes both augmented reality and affective artificial intelligence, and helps users learn key social and emotional skills.

**Method:** Twenty-one children and adults with ASD across a spectrum of severity used BPAS for a coaching session. The user’s tolerability to the smartglasses was determined through caregiver report, and user being able to wear the smartglasses for one-minute (initial tolerability threshold), and for the entire duration of the coaching session (whole session tolerability threshold).

**Results:** Nineteen out of 21 users (90.5%) demonstrated tolerability on all three measures. Caregivers reported 21 out of 21 users (100%) as tolerating the experience, while study staff found only 19 out of 21 users managed to demonstrate initial tolerability (90.5%). Of the 19 users who demonstrated initial tolerability, all 19 were able to use the smartglasses for the entire session (whole session tolerability threshold) (*n* = 19 of 19, 100%). Caregiver’s reported that 19 out of 21 users (90.5%) successfully used BPAS, and users surpassed their caregiver’s expectations in 15 of 21 cases (71.4%). Users who could communicate reported BPAS as being comfortable (94.4%).

**Discussion:** This preliminary report suggests that BPAS is well tolerated and usable to a diverse age- and severity-range of people with ASD. This is encouraging as these devices are being developed as assistive technologies for people with ASD. Further research should focus on improving smartglasses design and exploring their efficacy in helping with social communication in children and adults with ASD.

## Introduction

Modern smartglasses are small head-mounted displays that integrate a range of sensors that can capture video, audio, and movement data. Smartglasses can deliver an augmented reality (AR) experience, where the user can see virtual objects overlaid on top of their real-world view as they look through the optical display. Smartglasses delivering AR are believed to have considerable potential as educational and healthcare tools, and an increasingly wide range of smartglasses are available for developers and consumers (1, 2).

A wide range of assistive technologies have been developed for ASD and include smartphone and tablet apps, computer programs, social robots, and virtual reality (3). There have been encouraging findings about the positive impact of such technologies, yet many children and adults with ASD continue to have considerable unmet educational and healthcare needs. Interest has been growing in the use of AR as a teaching tool for children and adults with ASD, and understanding how people with ASD experience and are affected by head-mounted displays remain key questions that face the field (4). An AR experience can be delivered on a variety of different platforms including smartphones, tablets, stationary displays, and on “heads-up” smartglasses. Much of the current AR research has been on AR delivered through handheld/”heads-down” devices (5-7). Studies have demonstrated that AR delivered on smartphones, tablets, and desktop computers may help people with ASD with their attention (6), emotion recognition (8), ability to notice social cues (5), social skills (9), ability to engage in pretend play (10), and even as a navigation aid for planning trips (7). However, using AR is not a risk-free endeavor, children using smartphone-based AR have developed postural and grip strain, in addition to experiencing falls (11), and smartphone-based AR games can lead to injury through distraction, with resultant falls and major trauma already being reported (12).

Smartglasses may offer several advantages when compared to smartphone and tablet devices, and have been described as the platform of the future for AR (13). Use of smartglasses may be less distracting than smartphones, and may require less cognitive workload (14, 15). By looking through smartglasses, users can continue to look “heads-up” at the environment around them and also remain “hands-free”, because smartglasses are head-worn (16). These advantages may enable users to continue to observe the social world around them, something that is considerably impacted when using a smartphone (17). Additionally smartglasses allow users to keep their hands unoccupied, making it easier to use them in non-verbal communication and/or academic/occupational activities, which are particularly pertinent considerations for children and adults with ASD who demonstrate impairment in social communication (18). To our knowledge, we have published the first report of the feasibility of using AR smartglasses to provide social and cognitive coaching in children with ASD (16).

Research is required to determine the tolerability of AR smartglasses given that ASD is accompanied by a range of sensory, behavioral, and cognitive challenges that may make wearing such devices difficult. Many people with ASD have sensory sensitivities, and they may struggle to wear conventional prescription glasses (19), brush their teeth, or comb their hair (20, 21). Smartglasses often have a similar form factor to prescription glasses, and in the case of Glass Explorer Edition (formerly known as “Google Glass”), may weigh the same as typical pair of prescription lenses and frame (22). Unlike prescription glasses, smartglasses produce additional sensory stimulus in the form of visual input via their optical displays, and audio via their speakers. It is therefore important to study how people with ASD would respond to and tolerate wearing such devices. With the exception of conventional prescription glasses, there are only rare occasions when one would need to “wear” a face mounted object. In this regard, wearing smartglasses may be a particularly novel experience, with few daily life comparators. This is an important consideration because people with ASD can exhibit considerable distress when exposed to unfamiliar situations, changes in routine, or changes in environment (18). Despite the abovementioned concerns, there continues to be a dearth of research into AR smartglasses for people with ASD. The authors have found that many clinicians, educators, and people from the ASD community have expressed doubt as to whether children and adults with ASD would tolerate wearing AR smartglasses. This has led to the commonly encountered question, *but will they even wear it?* This is not surprising given that even wearing conventional glasses has been highlighted as a major challenge by prominent ASD charities (23).

The importance of understanding how people with ASD will respond to such devices is only heightened by understanding the potential benefits of conducting research with smartglasses. Smartglasses, like smartphones, contain a myriad of sensors, such as an accelerometer and camera, and are able collect video, audio, movement, physiologic and user interaction data (24). These quantitative data can be collected and analyzed in an effort to undertake digital phenotyping, and more importantly, to help support research efforts to help subtype highly clinically heterogeneous behavioral conditions such as ASD (25).

To explore the tolerability of AR smartglasses, we studied whether children and adults with ASD were able to tolerate wearing *the Brain Power Autism System* (BPAS), a novel social communication coaching smartglasses that utilizes AR and emotional artificial intelligence (16). BPAS has undergone feasibility (16), acceptability (26), safety (27), and clinical impact studies (28). BPAS is based on a highly-modified version of the Google Glass Explorer Edition (formerly Google, Mountain View, CA) and Glass Enterprise Edition (X, Mountain View, CA).

## Methods

The methods and procedures of this study were approved by Asentral, Inc., Institutional Review Board, an affiliate of the Commonwealth of Massachusetts Department of Public Health.

### User Recruitment

A sequential sample of 21 children and adults with clinically diagnosed ASD were recruited from a database of individuals who completed a web-based signup form expressing interest in participating in smartglasses research. Individuals represented a demographically and clinically diverse group comprised of different ages, genders, verbal abilities, and level of functioning (Table 1). Written consent was obtained from the legal guardians of children and from cognitively-able adults. Children between 7 – 17 years-old provided written assent when possible. In this report, every user was accompanied by a parent or other caregiver to the session, and users and caregivers could ask for the session to stop at any time and for any reason. All users completed the Social Communication Questionnaire as a means of understanding their level of social communication impairment (29). The Social Communication Questionnaire score demonstrates that the user sample represented a wide range of social communication impairment.

**Table 1:** Demographics.

| Table 1: Demographics |  |  |
| --- | --- | --- |
| Number of users | 21 |  |
| Mean age ± SD, years | 11.9 +/- 4.9 | Range = 4.4 years - 21.5 years |
| Participant gender | Male: 19 (90.5%) | Female: 2 (9.5%) |
| Verbal or nonverbal | Verbal: 19 (90.5%) | Nonverbal: 2 (9.5%) |
| Social Communication Questionnaire Score ± SD | 18.5 +/- 6.1 | Range = 6 - 28 |

### Data Collection Procedure

Users and caregivers were given an introductory explanation and demonstration of BPAS smartglasses. Users were then given the chance to wear the smartglasses (Figure 1), aided as needed by study staff and their caregivers for correct initial placement (Figure 2).

**Figure 1. (A-C).**
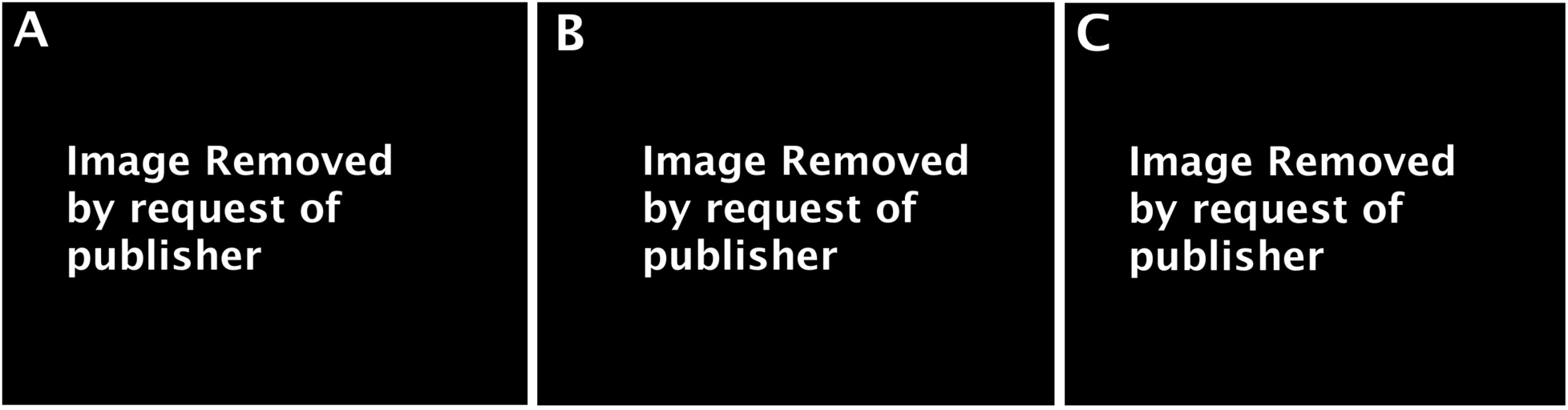
Three users with ASD wearing the Brain Power Autism System and using its socio-emotional coaching apps. *Pictures used with user / caregiver permission*.

**Figure 2. (A-B).**
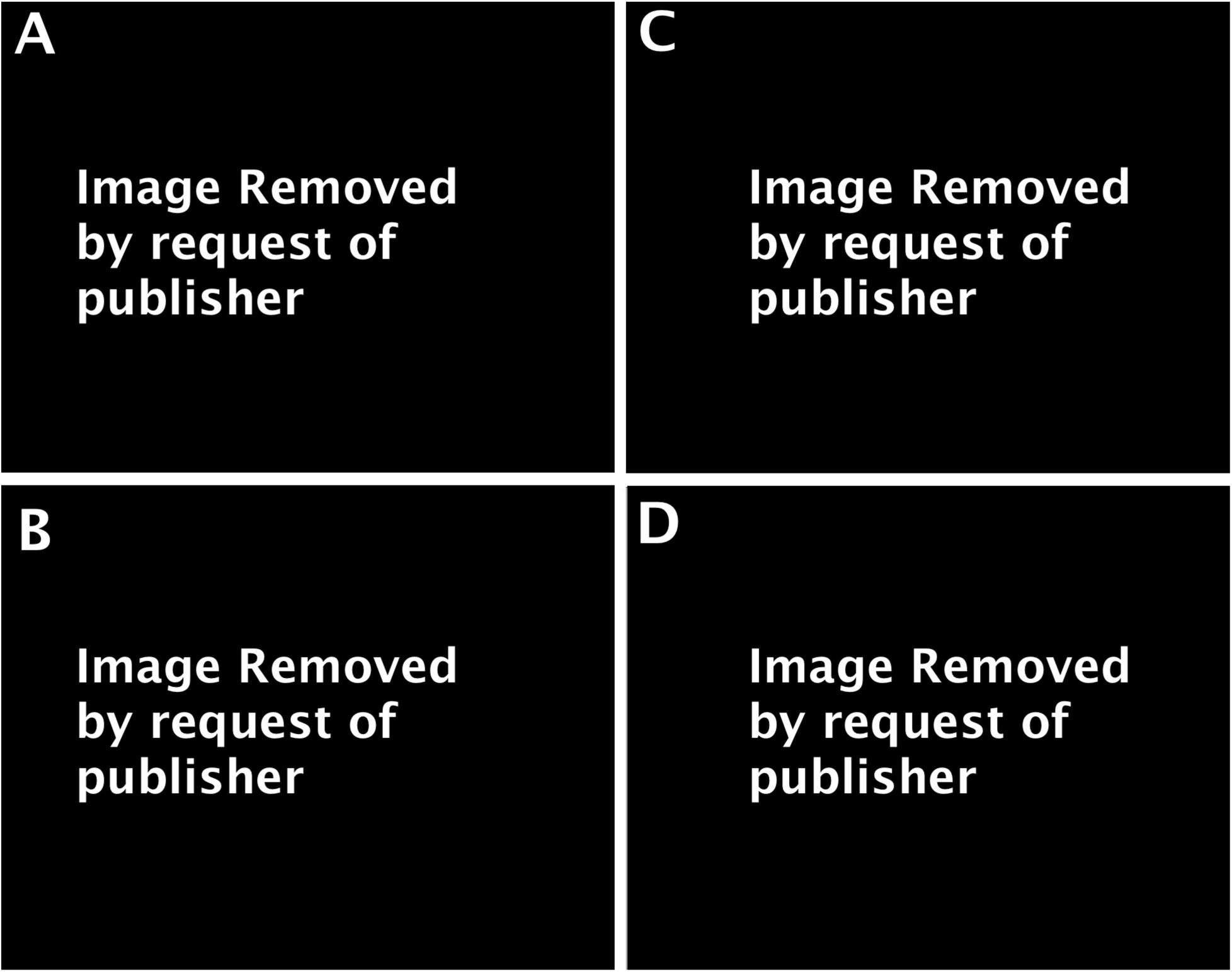
Caregiver assisting User to wear BPAS smartglasses during testing. User demonstrated tolerability on all 3 measures, and was witnessed to spontaneously hug his caregiver during use of social communication app use **(C, D)**. *Pictures used with user / caregiver permission*.

The user’s tolerability to the smartglasses was determined through caregiver report, the user’s ability to wear the smartglasses for one-minute (initial tolerability threshold), and the user’s ability to wear the smartglasses for the entire duration of the coaching session (whole session tolerability threshold). The initial tolerability threshold provides a rapid understanding of how well a user would respond to the physical form factor of the smartglasses, an important consideration given the unique set of sensory and cognitive challenges of each user. The whole session tolerability threshold represents how well the user tolerates wearing the smartglasses, but represents their use of the coaching apps as they undertake a series of structured activities with their caregiver lasting between 1-1.5 hours. At the end of the session, caregivers were able to rate how well they felt the user tolerated using BPAS through a five-point Likert scale (1 = very low, 5 = very high). A tolerability rating of low or very low as deemed to be a negative indication of tolerability, while neutral, high, or very high caregiver ratings were noted as an indication of tolerability.

Caregivers were also asked to use a five-point Likert scale to rate if they felt the user was able to successfully use BPAS with their assistance (1 = strongly disagree, 5 = strongly agree), and whether they felt the user responded more positively to BPAS smartglasses than the they had expected (1 = strongly disagree, 5 = strongly agree). For these responses, a higher standard had to be set compared to tolerability: a rating of agree/strongly agree (4 or 5) was determined to be a positive response for each of these questions. Users who could communicate verbally with their caregiver/study staff were asked to rate how comfortable the smartglasses were. Both caregivers and users were able to provide additional feedback to any question in the interviews.

## Results

Nineteen out of 21 users (90.5%) demonstrated tolerability on all three measures (caregiver report, initial tolerability threshold, and whole session tolerability threshold) (Table 2). Of the 19 users who managed to pass the initial tolerability threshold (*n* = 19 of 21, 90.5%), all went on to use BPAS for the entire coaching session, passing the whole session tolerability threshold (*n* = 19 of 19, 100%). Two users, both nonverbal, did not pass the initial one-minute tolerability threshold as they would not continue to wear the smartglasses once placed. Users who were verbal and able to answer questions (*n* = 18 of 21) rated the smartglasses as being comfortable to use (*n* = 17 of 18, 94.4%) (Table 3). A majority of caregivers felt users responded more positively to the smartglasses than they had expected (*n* = 15 of 21, 71.4%). A number of caregivers provided additional feedback, suggesting that users may benefit from extended and/or repeated orientation and introduction sessions with BPAS. The results are graphically represented in Figure 3.

**Table 2:** Tolerability report of BPAS.

| Table 2:<br>Tolerability report of BPAS | Yes | No |
| --- | --- | --- |
| Initial tolerability threshold (worn for at least 1 minute of continuous use) | 19/21,<br>90.5% | --- |
| Whole session tolerability threshold (worn until session completion) | 19/21,<br>90.5% | 2/21,<br>9.5% |
| Caregiver report of tolerability | 21/21,<br>100% | --- |
| Users demonstrating tolerability on all measures | 19/21,<br>90.5% | 2/21,<br>9.5% |

**Table 3:** User experience report.

| Table 3:<br>User experience report | Yes | Mixed response<br>/ Sometimes | No |
| --- | --- | --- | --- |
| User reported that experience was comfortable (only users who able to answer questions: $n = 18$) | 17/18,<br>94.4% | 1/18,<br>5.6% | 0/18,<br>0% |
| Caregiver reported that user managed to use the device with assistance | 19/21,<br>90.5% | --- | 2/21,<br>9.5% |
| Caregiver reported user responded more positively to the device than expected | 15/21,<br>71.4% | 6/21,<br>28.6% | 0/21,<br>0% |

**Figure 3:**
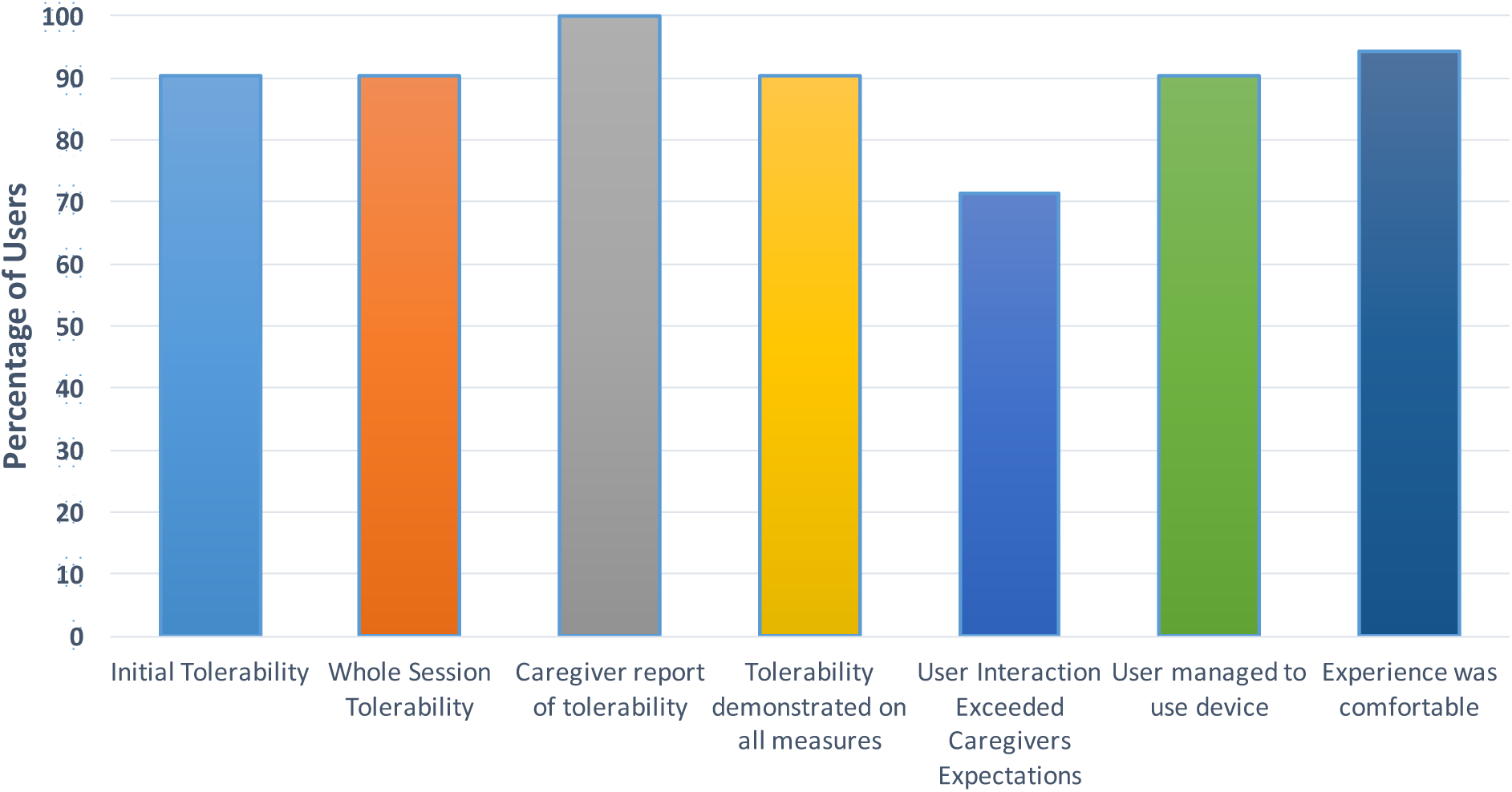
Summary of BPAS Tolerability and Experience.

## Discussion

Children and adults with ASD have considerable unmet educational and behavioral health needs, and technology-aided solutions may provide a scalable and effective tool to help address these demands for resources. While AR smartglasses have been designed as a social communication aid for people with ASD (16), there is only limited research to understand how acceptable and tolerable this technology is to individuals with ASD (16). Our preliminary study shows that a diverse range of children and adults with ASD can tolerate wearing and using the BPAS, AR smartglasses designed to function as a social communication aid for people with ASD.

Tolerability was demonstrated across all three of our measures in 90.4% of users. Every user who demonstrated initial one-minute tolerability managed to continue to demonstrate tolerability for the entire coaching session that ran between 1-1.5 hours. This suggests that the tolerability of users with ASD to such smartglasses can be accurately predicted based on their ability to initially use the device over a relatively short amount of time.

Our data help to answer our initial question, and title of the paper, but will they even *wear* it? In our experience this is one of the most common questions that parents, educators, and researchers ask us when BPAS is shown to them. These data show that children and adults with ASD can not only wear smartglasses for relatively lengthy durations of time, but are able to use them, and describe the experience as comfortable. How the users interacted with BPAS surpassed the expectations of their caregivers in a majority of cases. We did find that both non-verbal users struggled to wear the smartglasses, and were unable to pass the initial tolerability threshold. Based on feedback from caregivers, a more gradual introduction and orientation process to the smartglasses may have been useful in these cases.

Additionally, given that sensor-rich smartglasses are quantitative data gathering tools, it is helpful to know that they can be worn for such durations in people with behaviorally heterogeneous conditions that could benefit from digital phenotyping and subtyping.

While our results are promising, there are a number of limitations. Although this work is, to our knowledge, the first report of the tolerability of smartglasses as a social communication aid in people with ASD, our sample size is moderate (*n* = 21). Additionally, given the customized nature of BPAS, generalizability may be limited in the case of other smartglasses, different smartglasses software apps, and even an unmodified Google Glass device.

More longitudinal research would be useful to determine whether the tolerability that we have observed continues to last after repeated coaching sessions. Understanding the effects of repeated sessions over a longer duration of time is an important, as many training and educational programs for people with ASD involve repeated sessions over a long period of time. Further research is required to investigate the efficacy of AR smartglasses in ASD, but the tolerability and usability of such devices does not appear to be a substantial barrier to their use.

## References

1. Elder S, Vakaloudis A, editors. A technical evaluation of devices for smart glasses applications. Internet Technologies and Applications (ITA), 2015; 2015: IEEE.

2. Elder S, Vakaloudis A, editors. Towards uniformity for smart glasses devices: An assessment of function as the driver for standardisation. Technology and Society (ISTAS), 2015 IEEE International Symposium on; 2015: IEEE.

3. Grynszpan O, Weiss PL, Perez-Diaz F, Gal E. Innovative technology-based interventions for autism spectrum disorders: a meta-analysis. Autism. 2014;18(4):346-61.

4. Newbutt N, Sung C, Kuo HJ, Leahy MJ. The acceptance, challenges, and future applications of wearable technology and virtual reality to support people with autism spectrum disorders. Recent Advances in Technologies for Inclusive Well-Being: Springer; 2017. p. 221-41.

5. Chen C-H, Lee I-J, Lin L-Y. Augmented reality-based video-modeling storybook of nonverbal facial cues for children with autism spectrum disorder to improve their perceptions and judgments of facial expressions and emotions. Computers in Human Behavior. 2016;55:477-85.

6. Escobedo L, Tentori M, Quintana E, Favela J, Garcia-Rosas D. Using augmented reality to help children with autism stay focused. IEEE Pervasive Computing. 2014;13(1):38-46.

7. McMahon D, Cihak DF, Wright R. Augmented reality as a navigation tool to employment opportunities for postsecondary education students with intellectual disabilities and autism. Journal of Research on Technology in Education. 2015;47(3):157-72.

8. hen C-H, Lee I-J, Lin L-Y. Augmented reality-based self-facial modeling to promote the emotional expression and social skills of adolescents with autism spectrum disorders. Research in developmental disabilities. 2015;36:396–403.

9. Tentori M, Hayes GR, editors. Designing for interaction immediacy to enhance social skills of children with autism. Proceedings of the 12th ACM international conference on Ubiquitous computing; 2010: ACM.

10. Bai Z, Blackwell AF, Coulouris G. Using augmented reality to elicit pretend play for children with autism. IEEE transactions on visualization and computer graphics. 2015;21(5):598-610.

11. Radu I, Guzdial K, Avram S, editors. An Observational Coding Scheme for Detecting Children’s Usability Problems in Augmented Reality. Proceedings of the 2017 Conference on Interaction Design and Children; 2017: ACM.

12. Franchina M, Sinkar S, Ham B, Lam GC. A blinding eye injury caused by chasing Pokémon. The Medical Journal of Australia. 2017;206(9):384.

13. Monkman H, Kushniruk AW, editors. A see through future: augmented reality and health information systems. ITCH; 2015.

14. He J, Choi W, McCarley JS, Chaparro BS, Wang C. Texting while driving using Google Glass™: Promising but not distraction-free. Accident Analysis & Prevention. 2015;81:218–29.

15. He J, Ellis J, Choi W, Wang P. Driving while reading using Google glass versus using a smart phone: which is more distracting to driving performance. City. 2015.

16. Liu R, Salisbury JP, Vahabzadeh A, Sahin NT. Feasibility of an Autism-Focused Augmented Reality Smartglasses System for Social Communication and Behavioral Coaching. Frontiers in Pediatrics. 2017;5(145).

17. Chen P-L, Pai C-W. Smartphone gaming is associated with pedestrians′ head-turning performances: an observational study of street-crossing behaviours at uncontrolled intersection in Taipei. International Journal of Sustainable Transportation. 2017(just-accepted):00-.

18. Association AP. Diagnostic and Statistical Manual of Mental Disorders (DSM-5®): American Psychiatric Pub; 2013. But will they even wear it? Exploring the tolerability of social communication coaching smartglasses in children and adults with autism.

19. DeLeon IG, Hagopian LP, Rodriguez-Catter V, Bowman LG, Long ES, Boelter EW. Increasing wearing of prescription glasses in individuals with mental retardation. J Appl Behav Anal. 2008;41(1):137-42.

20. Smith MD, Belcher R. Teaching life skills to adults disabled by autism. Journal of Autism and Developmental Disorders. 1985;15(2):163-75.

21. Matson JL, Taras ME, Sevin JA, Love SR, Fridley D. Teaching self-help skills to autistic and mentally retarded children. Research in Developmental Disabilities. 1990;11(4):361-78.

22. Walsh G. The weight of spectacle frames and the area of their nose pads. Ophthalmic and Physiological Optics. 2010;30(4):402-4.

23. Speaks A. Autism Challenge for our Experts: Kid Needs Glasses; Won’t Tolerate Anything on Face.: Autism Speaks; 2012

24. Hernandez J, Li Y, Rehg JM, Picard RW, editors. Bioglass: Physiological parameter estimation using a head-mounted wearable device. Wireless Mobile Communication and Healthcare (Mobihealth), 2014 EAI 4th International Conference on; 2014: IEEE.

25. Onnela JP, Rauch SL. Harnessing Smartphone-Based Digital Phenotyping to Enhance Behavioral and Mental Health. Neuropsychopharmacology. 2016;41(7):1691-6.

26. Sahin NT, Keshav NU, Salisbury JP, Vahabzadeh A. Cool Enough for School: Second Version of Google Glass Rated by Children Facing Challenges to Social Integration as Desirable to Wear at School. bioRxiv. 2017.

27. Sahin NT, Keshav NU, Salisbury JP, Vahabzadeh A. An Augmented Reality Social Communication Aid for Children and Adults with Autism: User and caregiver report of safety and lack of negative effects. bioRxiv. 2017.

28. Vahabzadeh A, Keshav NU, Salisbury JP, Sahin NT. Preliminary Report on the Impact of Smartglassesbased Behavioral and Social Communication Aid on Hyperactivity in Children and Adults with Autism. bioRxiv. 2017.

29. Rutter M, Bailey A, Lord C. The social communication questionnaire: Manual: Western Psychological Services; 2003.

